# The role of RNA in the nanoscale organization of alpha-synuclein phase separation

**DOI:** 10.1101/2024.09.03.610813

**Authors:** Sabrina Zappone, Eleonora Perego, Jakob Rupert, Elsa Zacco, Eli Slenders, Gian Gaetano Tartaglia, Giuseppe Vicidomini

## Abstract

The cellular accumulation of alpha-synuclein (aS) aggregates is a hallmark of several neurodegenerative diseases. Recent studies suggest that the aberrant transition of monomeric aS into solid-like aggregates may occur through an intermediate liquid-like state, where the protein partitions between dense and dilute phases. Although aS is not typically recognized as an RNA-binding protein, it can bind RNA under aggregation conditions, but its impact on aS liquid-like phases remains unexplored. Employing a combination of fluorescence spectroscopy techniques, we investigated aS dynamics in both phases in the presence of RNA. Our analysis revealed the formation of nanoclusters involved in initiating phase separation and uncovered heterogeneity within the dense phase, discovering that aS molecules exist in two distinct mobility states. Additionally, we demonstrated that RNA induces morphological changes and promotes the liquid-to-solid transition of aS dense phase. These findings underscore the active role of RNA in modulating aS phase transitions.

## Introduction

Protein misfolding, aggregation, and its associated loss of function have long been associated with several human disorders. Misfolded proteins usually convert into fibrillar aggregates known as amyloids (1). While the amyloidogenesis of aggregation-prone proteins, such as amyloid beta and alpha-synuclein, has been extensively studied and characterized *in vitro* (2–4), its mechanisms in the cellular context have initially remained less clear. In recent years, the process of protein aggregation has been continuously more linked to the process of liquid-liquid and liquid-to-solid phase transitions (5). Liquid-liquid phase separation (LLPS) is a thermodynamic-driven process in which biomolecules, by establishing weak intermolecular interactions, self-organize into a liquid-like dense phase, immersed into another liquid-like dilute phase (4). The accumulation of molecules in the dense phase takes form into micron-sized condensates termed membrane-less organelles (MLOs). MLOs have been shown to be critical for many cellular functions as a way of tuning reactions in space and time, sequestering toxic molecules, or organizing cellular content (6).

Recent progress has also shown that the ability to undergo LLPS is not exclusive to the components of MLOs, but occurs at a wide range in the cellular proteome (7) and a much lower scale than the micrometer scale (8). The phase separation of proteins is triggered only when molecules reach a certain concentration, called saturation concentration (C_sat_). Above C_sat_, molecules follow the classical nucleation theory, which implies the formation of a system composed of a homogeneous dilute phase coexisting with a micron-sized dense phase (henceforth, macro-condensates). The formation of any molecular complex that is different than macro-condensates is energetically penalized. However, it was recently demonstrated that some phase-separating proteins deviate from the classical nucleation theory by forming nanometer-sized clusters, immersed in the dilute phase, both below and above C_sat_ (8). Nanoscale condensates, also termed nanoclusters, have been observed for many different phase-separating proteins and have been shown to be primarily involved in the early nucleation processes of crystals, condensates, and aggregate growth (9).

One particular case in which nanoclusters have been characterized as a critical state-of-matter is the protein alphasynuclein (aS) (10). aS is an intrinsically disordered protein found abundantly expressed in mammalian cells (11). It has been associated with numerous cellular processes, including synaptic vesicle trafficking, exocytosis, and DNA repair (11– 13). aS has sparked interest in the scientific community for its involvement in a number of neurodegenerative diseases collectively known as synucleopathies (14). The transition of aS from a soluble, monomeric state to an insoluble, irreversible state results in the accumulation of amyloid aggregates within cells. This phenomenon, known as liquid-to-solid transition, has long been accepted as the pathological pathway behind the deposition of aS monomers into amyloid aggregates (15). However, the LLPS and the following transition of liquid-like condensates to solid-like ones have been recently proposed as an alternative pathway (16). Indeed, the high local concentration and confinement of molecules inside liquid-like condensates can stimulate the primary nucleation and the formation of solid aggregates at physiological pH levels, a process further exacerbated by the potential uptake of pre-aggregated forms of aS (17, 18). As mentioned, aS also rapidly forms nanoclusters in solution that persist even in the presence of macro-condensates, further indicating LLPS as a potential mechanism of aggregate formation and propagation (10). Even though the regulation or disruption of phase separation of certain proteins can occur at the level of nanoclusters (8), their impact on aS LLPS has not been investigated yet.

Apart from the formation of nanoclusters, aS phase transitions can also be influenced by interactions with other partners, both proteins and nucleic acids (19, 20). Despite the total amount of protein, the concentration of aS within condensates is estimated to be three orders of magnitude higher than the dilute phase (17). These exceedingly high concentrations kinetically facilitate specific interactions that appear transient under solution conditions, such as interactions with RNA. Besides being a protein-mediated process, RNA especially has been shown to be one of the critical players in the initiation and regulation of protein phase separation (21, 22). aS is not known to bind RNA in the monomeric state but has been shown to exhibit a large increase in RNA-binding propensity upon aggregation and in the ability to sequester RNA in its aggregated forms (23). The effect of RNA on aS phase transition has been far from clear and studied primarily with polynucleotides, such as poly(U) and poly(A): poly(U) molecules have been shown to ameliorate aS aggregation (24), while poly(A) promotes the localization of aS on the condensate surface and accelerates its aggregation (25). Nevertheless, there is no evidence on how RNA can influence the LLPS of aS both at nano- and micro-scales, nor its consequential impact on pathological liquid-to-solid phase transition.

The investigation of LLPS systems has been predominantly focused on the dense phase. However, characterizing the dilute phase is fundamental for achieving a comprehensive understanding of phase-separated systems. This task poses significant technical challenges due to the inherent heterogeneity of the dilute phase and the potential formation of nanoclusters. Phase-separated systems have been conventionally studied using several methods, such as mass spectroscopy (26), dynamic light scattering (27), nuclear magnetic resonance spectroscopy (28), and fluorescence microscopy techniques. While fluorescence confocal microscopy imaging has been extensively used for visualizing macro-condensates, it suffers from poor temporal resolution (in terms of frame rate) and spatial resolution for quantifying the dynamics of sub-diffraction-limited species such as nanoclusters. Fluorescence recovery after photobleaching (FRAP), typically implemented in a confocal laser scanning microscope, is widely used for characterizing intra-condensate dynamics. However, FRAP measurements are often difficult to interpret as they depend on the optical system, laser power, and sample geometry (29). They are mostly suitable for investigating the immobile fraction or very slow-moving molecules, making FRAP ineffective for studying the fast dynamics in the dilute phase. Phase-separating proteins like aS show diverse temporal dynamics, with very fast movement in the dilute phase (nanoclusters and monomers) and slower movement in the dense phase. As a result, relying solely on FRAP does not reveal all the underlying phenomena.

These limitations can partially be overcome with fluorescence fluctuation-based methods. Among them, fluorescence correlation spectroscopy (FCS) measures the mobility of the molecules by statistically analyzing the temporal correlation of fluctuations in fluorescence intensity (30). To retrieve diffusion properties with FCS, molecules must move within a well-defined observation volume, such as one created by focusing a laser beam in a confocal microscope. FCS has already been applied to various LLPS scenarios, including identifying nanoclusters in the phase-separated system of pyrenoid (31) and amyloid (32) proteins. Both cases demonstrated that, due to its high sensitivity, FCS is a suitable method for studying molecular interactions in nano- and macro-condensates.

While valuable for characterizing molecular species based on their different mobility, conventional FCS cannot distinguish the spatial heterogeneity inherent in LLPS systems. Spot-variation FCS (svFCS) addresses this limitation by performing FCS measurements with increasing observation volumes. This FCS variant enables the determination of not only diffusion coefficients but also of the type of movement in the sample, thereby resolving the heterogeneity of the system (33). Here, we enhance conventional confocal FCS measurements by replacing the typical single-element detector of confocal microscopy, such as a photomultiplier or a single-photon avalanche diode (SPAD), with a SPAD array detector (34). The array detector preserves the spatial distribution of the fluorescent signal of the sample, which would otherwise be lost with a single-element detector. This spatial information allows for a straightforward and instantaneous implementation of svFCS (35). Moreover, the single-photon timing capability of the SPAD facilitates photon time-tagging detection (36), which allows for sampling of the signal photon-by-photon and with high temporal precision (down to the sub-nanosecond scale). This high temporal resolution allows for the resolution of fluorescence lifetime changes. In the context of LLPS, fluorescence lifetime has proven valuable in distinguishing molecular packing densities of proteins such as Tau (37) and G3BP1 (38) within MLOs.

We have already shown that this suite, referred as photon-resolved microscopy, facilitates analysis across spatial scales from micrometers to nanometers and temporal scales from seconds to nanoseconds, enabling multiplexed analysis of condensate dynamics with a single measurement (38). The simultaneity and relatively short duration (tens of seconds) of the measurements guarantee reduced complexity, lower phototoxicity, while preserving temporal heterogeneity.

By employing this platform, we characterized the heterogeneity of aS dynamics across different temporal scales, identified the nanocluster formation, and discriminated freely diffusing and domain-constrained molecules both in and out of macro-condensates. We have also successfully investigated the effect of RNA on the aS phase transition. Strikingly, our results show that the presence of RNA drastically alters the shape and the internal dynamics of aS condensates towards a solid-like state. These findings further suggest that RNA acts as a strong modifier of aS aggregation, by influencing not only the direct prion-like conversion into an amyloid state (23), but also the liquid-liquid and liquid-solid phase transitions of aS condensates.

## Results

### *In vitro* reconstitution of alpha-synuclein phase separation

We first characterized the dynamics of purified full-length alpha-synuclein (aS) by fluorescence correlation spectroscopy (FCS) under non-LLPS-inducing conditions (figure 1, 2A top). A one-diffusing component model was sufficient to fit the FCS autocorrelation data, hinting at a uniform, non-interacting population of freely diffusing molecules (figure S2). The retrieved diffusion coefficient, *D*_app_ = 134 ± 4 µm^2^/s, and its related hydrodynamic radius, *r*_H_ = 1.65 ± 0.05 nm (calculated with the Stokes-Einstein formalism), are both compatible with literature values for monomeric aS (39– 42).

**Fig 1.**
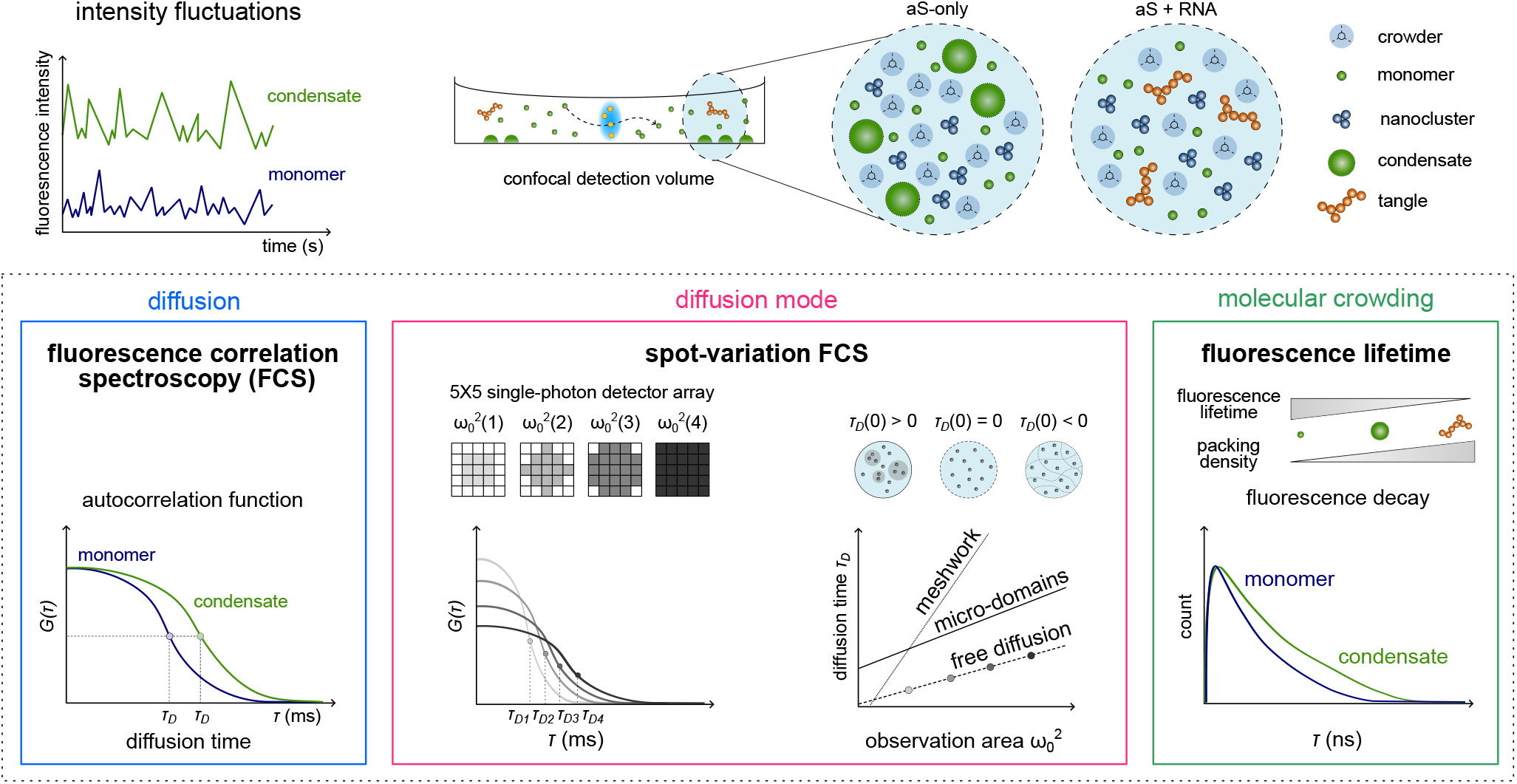
Schematic representation of the experimental and analysis session. The phase separation assay is a mixture of aS, aS-atto488, and, in the case of the RNA experiment, total yeast RNA. The sample is placed into a closed chamber and under a confocal laser scanning microscope equipped with a 5×5 single-photon array detector. Fluorescence over time is acquired in different sample positions. The same dataset of fluorescence over time can be used to perform fluorescence lifetime quantification or fluorescence correlation spectroscopy-based methods (FCS and spot-variation FCS).

**Fig 2.**
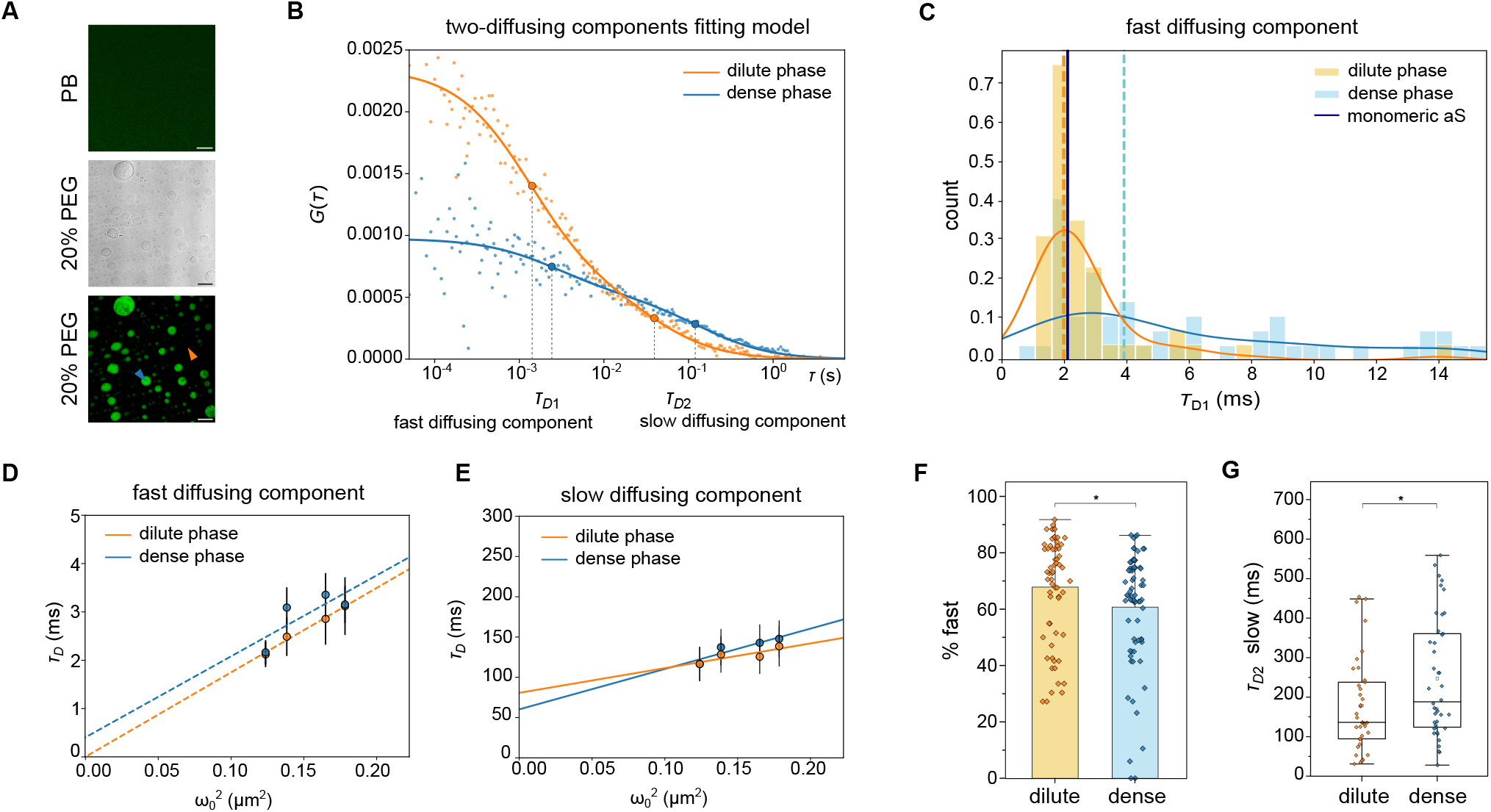
aS forms nano- and micro-complexes in a phase-separated system. (A) Confocal image (top) of 700 nM monomeric aS-atto488 in PB. Phase contrast microscopy image (middle) and confocal (bottom) image of aS in LLPS buffer. In the presence of 20% PEG, aS spontaneously forms a dense phase (blue arrow) surrounded by a dilute phase (orange arrow). Scale bars: 5 µm. (B) Representative autocorrelation functions in the dilute (in orange) and the dense (in blue) phase of aS, as indicated by the arrows in (A). The lines represent the fit of the autocorrelation curves with a two-components fitting model. Diffusion times for the fast (*τ* _*D*1_) and slow (*τ* _*D*2_) diffusing components are shown. (C) Histogram of diffusion times *τ* _*D*1_ measured for the fast diffusing component in the dense (blue) and dilute (orange) phases. The solid lines represent the upper envelope of the histograms. The dashed lines represent the median of both distributions; the vertical blue solid line indicates the median of the *τ* _*D*_ for monomeric aS in the LLPS buffer. *n* = 50 (dilute phase), 53 (dense phase). (D, E) svFCS analysis of the fast (D) and slow (E) diffusing component for both dilute and dense phases. Diffusion times are plotted against the observation areas *ω*_0_^2^ to retrieve the diffusion law *τ* _*D*_ (*ω*_0_^2^) (dashed line). Data points and error bars are, respectively, mean and standard deviations of multiple measurements (*n* = 4 independent experiments). (F) Percentage of the fast diffusing component in the dilute and the dense phases. *n* = 40 (dilute phase), 48 (dense phase). Mann-Whitney statistical test (*: p-value *<* 0.05). (G) Box-plot of diffusion times *τ* _*D2*_ measured for the slow diffusing component in the dilute and the dense phase. *n* = 37 (dilute phase), 42 (dense phase). Mann-Whitney statistical test (*: p-value *<* 0.05). The horizontal line in each box represents the median, the edges are the 25th and 75th percentile, and the vertical line extends to the minimum and maximum data points.

Next, we measured the effect of the molecular crowder polyethylene glycol (PEG-8000) on aS liquid-liquid phase separation (LLPS) at two different concentrations (10% and 20%). In the presence of 10% PEG, the protein at the concentration of 50 µM has been reported not to undergo immediate and spontaneous macroscopic phase separation, while at the higher concentrations (20% PEG) it is supposed to form liquid-like condensates (10). As expected, confocal images do not reveal the formation of a visible dense phase for measurements at 10% PEG (figure S3B). However, FCS measurements still detect multiple diffusing components. Upon the addition of PEG, the autocorrelation function *G*(*τ*) could not be described anymore by a one-diffusing component model only (figure S5A). The characteristic diffusion time *τ* _*D*_ of the autocorrelation curve (the time at which the correlation amplitude halves) is directly associated with the diffusion coefficient and, consequently, the dynamics of the sample. A longer *τ* _*D*_ suggests slower dynamics, while a shorter *τ* _*D*_ indicates faster dynamics. By employing a two-diffusing component model, we detected the presence of a fast diffusing component with *τ* _*D*_ = 0.68 ± 0.04 ms and a slow diffusing one with *τ* _*D*_ = 95 ± 9 ms (figure S5A).

Further increasing the concentration of PEG to 20% triggers macroscopic LLPS, which is confirmed by the formation of spherical, micron-sized condensates (figure 2A, middle and bottom). To characterize the dynamics of aS at the nanoscale, we performed FCS measurements in the dilute and dense phases (figure 2B, figure S5B-C). Compared to the dilute phase, the autocorrelation curve *G*(*τ*) of the dense phase shows a smaller amplitude *G*(0) and a shift towards longer time delays. The amplitude of the autocorrelation curve is inherently linked to the average number of molecules within the observation volume (Supplementary Information), indicating a higher concentration of molecules in the dense phase. The observed shift of *G*(*τ*) of the dense phase towards longer time delays suggests overall slower dynamics than the dilute phase, potentially indicative of protein oligomerization or viscosity increase (figure 2B-G).

As for 10% PEG, two distinct diffusing components are detected in the FCS curves for both phases (figure S5B-C). To characterize them further, we analyzed their diffusion modes by performing spot-variation FCS (svFCS). If the molecule moves with Brownian motion, the diffusion time will decrease or increase correspondingly as the observation area changes. If the diffusion time *τ* _*D*_ does not increase or decrease proportionally with the change in the observation area 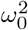, the type of movement in the sample is no longer described by free motion. By plotting the diffusion law 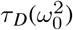 (the diffusion time versus the observation area), we can evaluate the type of movement with linear regression: if the intercept of the linear regression *τ* _D_(0) is 0, molecules are freely diffusing; if negative, molecules are highly restricted in mobility (such as inside a dense meshwork of cytoskeleton); if positive, molecules are moving in a domain-confined sample (33) (figure 1). In the latter, the intercept value is directly proportional to the confinement strength, which reflects the energy barrier needed for the molecules to exit the microdomain (43). The analysis revealed a free diffusion mode for the fast diffusing component (*τ* _*D*_(0) = 0.003 ± 0.214 ms for the dilute phase, *τ* _*D*_(0) = 0.4 ± 1.4 ms for the dense phase) (figure 2D). We hypothesized that the fast component refers to monomeric aS, freely diffusing in both dilute and dense phases. To validate our hypothesis, we analyzed the distribution of the diffusion times *τ* _*D*_ of the fast diffusing component in both phases (figure 2C). Overall, the diffusion time *τ* _*D*_ can be influenced by changes in viscosity and/or hydrodynamic radius (Supplementary Information). Assuming that in the dilute phase the viscosity (20% PEG buffer) is constant over time, *τ* _*D*_ is mainly affected by changes in the hydrodynamic radius and, ultimately, in the mono-or oligo-meric state of aS. To estimate the diffusion time of the monomeric aS in LLPS buffer, we measured the diffusion time of GFP both in phosphate buffer (PB) and 20% PEG buffer (figure S4). Since GFP does not phase separate or oligomerize, potential changes in the diffusion time are exclusively due to a difference in the buffer viscosity (figure S3A). Comparing *τ* _*D*_ of GFP measured in both buffers, we retrieved the viscosity of the 20% PEG buffer and, ultimately, the expected *τ* _*D*_ of monomeric aS (*τ* _*D*_ = 2.13 ± 0.07 ms, figure S4). The median of the *τ* _*D*_ distribution of the dilute phase (figure 2C, *τ* _*D*_ = 1.80 ms) corresponds well to the *τ* _*D*_ associated with the monomeric protein, confirming that aS is mainly organized in a freely diffusing monomeric state within the dilute phase. On the other hand, the heterogeneous distribution of *τ* _*D*_ for the dense phase reflects a more complex micro-environment (figure 2C). Finally, we compared the percentage of the fast diffusing component in the dense and dilute phases. We measured a higher percentage of the aS fast diffusing component (figure 2F) for the dilute phase, hinting again at a more complex micro-environment inside the condensates.

Regarding the slow diffusing component, svFCS revealed micro-domain confinement for both the dense and the dilute phase, as, in both, the intercept of the linear regression of the diffusion law 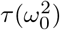 is bigger than zero (*τ* _*D*_(0) = 81 ± 22 ms for the dilute phase, *τ* _*D*_(0) = 60 ± 25 ms for the dense phase, figure 2E). While the diffusion of some molecules seems limited, liquid-like condensates are maintained in a dynamic equilibrium that allows constant exchange of molecules between the two phases. Considering this, we speculate that domain-confined aS in the dense phase reflects the portion of aS forming weak intermolecular interactions, which are essential for condensate formation. In contrast, the freely diffusing portion is the one readily exchanged with the external environment.

Interestingly, a domain-confined component is also present in the dilute phase. Mass photometric studies on alphasynuclein phase-separated systems proposed the presence of nanoscale assemblies within the dense phase. The domain-confined component we detected has all the properties attributed to nanoclusters: (i) the formation is immediate and spontaneous; (ii) the assembly also occurs upon subsaturated concentrations, where no instantaneous macroscopic LLPS is triggered (10% PEG buffer); (iii) aS does not show free diffusion within nanoclusters (figure 2E) (10). These observations allow us to associate the slow diffusing component in the dilute phase with nanoclusters.

Comparing the slow diffusing component in both phases, the diffusion time *τ* _*D*_ and the intercept value *τ* _*D*_(0) show an opposite trend. While *τ* _*D*_ of nanoclusters is lower than the one in macro-condensates (figure 2G), the intercept *τ* _*D*_(0) is bigger (figure 2E). A lower size of the complexes can explain the lower *τ* _*D*_ measured for nanoclusters. At the same time, being *τ* _*D*_(0) proportional to the confinement strength (43), we can infer that molecules in nanoclusters have a reduced mobility than the ones in macro-condensates.

### RNA does not affect the alpha-synuclein dilute phase

Once we had characterized the dynamics of aS under LLPS conditions, we studied the potential effect of RNA on the same *in vitro* system. In this work, we used total yeast RNA (henceforth referred to as RNA), which is known to undergo phase transitions in a PEG-based buffer. Depending on the concentration and the molecular weight of PEG, total RNA undergoes phase separation, forming either liquid-like, round-shaped condensates or irregular-shaped structures termed tangles (44). To confirm the behavior of RNA in LLPS buffer (20% PEG), we tested three different concentrations of RNA (5, 50, and 500 ng µL) labeled with SYTO™13 under the same LLPS conditions used for aS. By confocal imaging, we confirm the spontaneous formation of spherical condensates at all RNA concentrations (figure 3A, first column).

**Fig 3.**
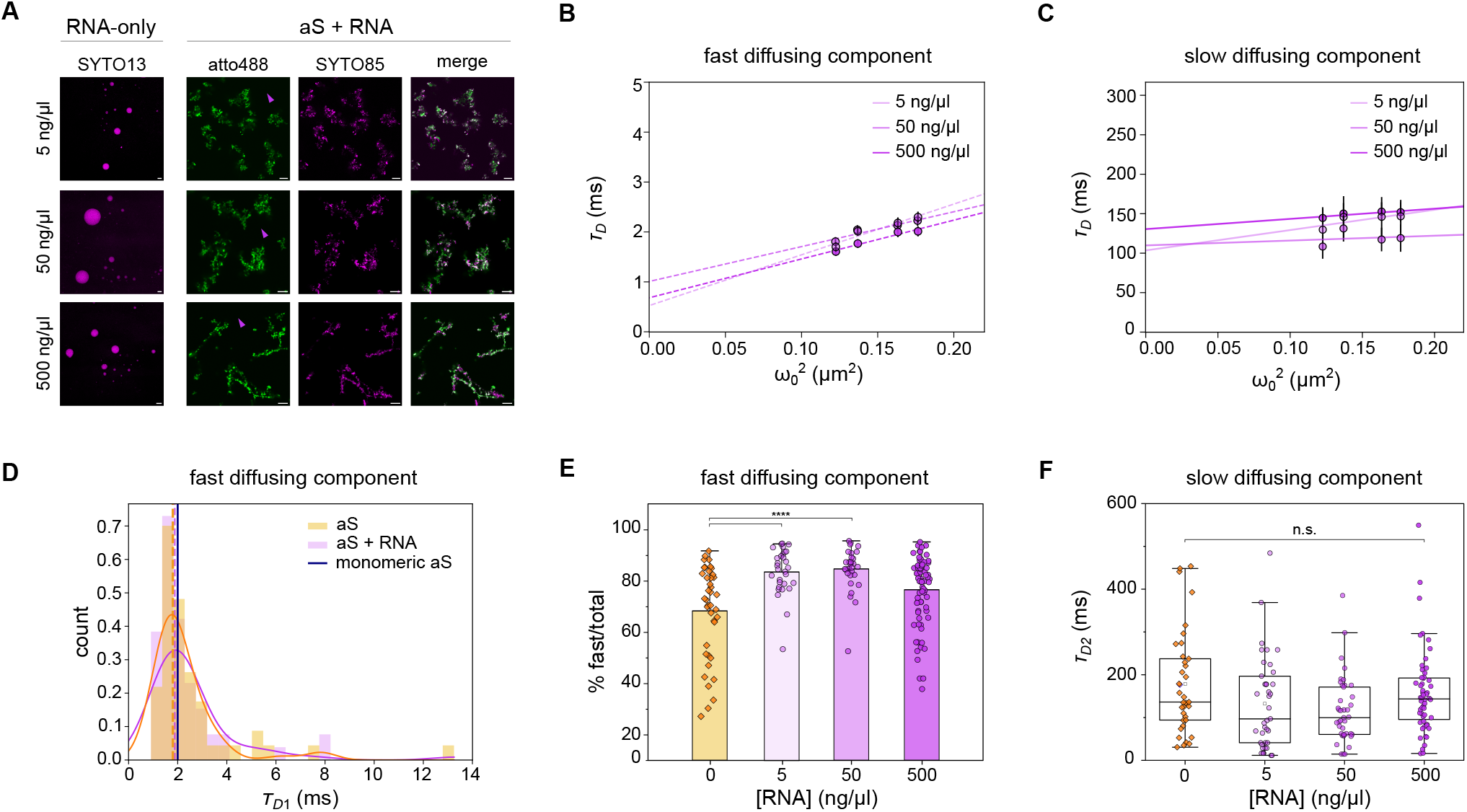
Effect of total yeast RNA on aS phase-separated system. (A) On the left, confocal images of 5, 50, and 500 ng µL RNA in LLPS buffer. On the right, confocal images of aS, aS-atto488 and RNA at different concentrations of RNA. A merge of both channels is shown on the right. Scale bars: 5 µm. (B, C) svFCS analysis of the fast (B) and slow (C) diffusing components of aS in the presence of RNA within the dilute phase. Diffusion times *τ* _*D*_ are plotted against the observation areas *ω*_0_^2^. Data points and error bars are, respectively, mean and standard deviations of multiple measurements (*n* = 4 independent experiments). (D) Histogram of diffusion times *τ* _D_ for the fast diffusing component of aS in the dilute phase under LLPS condition alone (orange) and in the presence of RNA (500 ng µL, purple). The solid lines on top of the histograms represent the upper envelope of the histogram. The dashed lines represent the medians of both distributions (1.88 ms for RNA); the vertical blue solid line indicates the median of the *τ* _*D*_ measured for the monomeric aS. *n* = 50 (aS), 57 (aS + RNA). (E) Percentage of the fast diffusing component in the dilute phase of aS in the presence of RNA. From left to right, *n* = 40, 30, 31, 77. Kruskal-Wallis statistical test followed by Dunn’s test (****: p-value *<* 0.00005). (F) Box plots of the slow diffusion times *τ* _D2_ in the dilute phase of aS alone and in the presence of RNA. No significant difference is appreciated among all conditions. From left to right, *n* = 38, 39, 41, 56. Kruskal-Wallis statistical test (n.s.: p-value *>* 0.05).

Next, aS and RNA were combined, and phase separation was triggered in the LLPS buffer. When LLPS of aS is induced in the presence of RNA, the dense phase shows an irregular shape rather than the spherical one obtained for RNA-or protein-only condensates (figure 2A, figure 3A, figure S3C). Due to surface tension, liquid-like condensates have a spherical shape. Since aggregated aS is packed in irregular-shaped and fibrillar structures, circularity is an ideal parameter for discriminating between liquid-like and solid-like condensates (45–47). We analyzed several confocal images to quantify the circularity (4*π*(area/perimeter^2^)) of condensates in the presence or absence of total RNA (figure S8A). We observed a strong reduction of the parameter (1 indicates a perfect sphere, 0 indicates a more elongated shape) upon adding RNA (figure S8B). The loss in the circularity and the structural transition towards an irregular shape suggests that aS, in the presence of total RNA, phase separates into tangles. Interestingly, these structures were never observed for the aS-only condensates. Moreover, no visible concentration-dependent effect regarding the size and number of tangles can be appreciated by confocal images (figure 3A). The staining of the RNA with the nucleic acid dye SYTO™85 also revealed the co-localization of RNA with the protein, confirming the actual recruitment of the nucleic acid within the dense phase (figure 3A).

We performed svFCS in both dilute and dense phases to explore the effect of RNA on aS LLPS at the nanoscale. As an initial step, we ruled out any potential changes in the viscosity of the LLPS buffer caused by the addition of RNA, as this could affect *τ* _*D*_ and thus future considerations regarding the molecular organization of aS in the dilute phase (figure S6). Measurements were acquired at all three previously tested RNA concentrations to assess any potential concentration-dependent effects on molecular mobility. As for the protein-only LLPS system, conventional FCS measurements within the dilute phase reveal the presence of two diffusing components (figure S7A-B-C). Regarding the fast diffusing component in the dilute phase, svFCS analysis indicated that aS freely diffuses in the sample (*τ* _*D*_(0) = 0.52 ± 0.30, 1.01 ± 0.25, 0.68 ± 0.18 ms, respectively for 5, 50, and 500 ng µL RNA, figure 3B). The distribution of the diffusion times *τ* _*D*_ associated with the fast component shows that the freely diffusing protein within the dilute phase is mainly monomeric. Additionally, the median of the distributions is comparable with the one measured for aS-only dilute phase (figure 3D for 500 ng/µL RNA, figure S7D for 5, 50 ng µL RNA). Possibly, this component represents the monomeric aS not recruited by the condensates yet.

The slow diffusing component in the dilute phase exhibits micro-domain confinement as diffusion mode across all tested RNA concentrations (*τ* _*D*_(0) = 103 ± 23, 110 ± 40, 130 ± 8 ms respectively for 5, 50, and 500 ng µL RNA, figure 3C). Detecting a slow diffusing component within the dilute phase provides evidence that nanocluster formation occurs even in the presence of RNA. No significant differences in confinement strength *τ* _*D*_(0) are observed among the three RNA concentrations. Although the presence of RNA does not affect the diffusion time (and thereby, the oligomerization) of the nanoclusters (figure 3F), an increase in confinement strength is observed.

Surprisingly, while no differences are seen in the dilute phase in terms of dynamics when RNA is present, the average percentage of the fast diffusing component is higher at low RNA concentrations (5 and 50 ng µL). This effect disappears when RNA concentration is increased up to 500 ng/µL. Indeed, at the highest tested RNA concentration, the aS dynamics do not change compared to the protein-only condition (figure 3E).

### RNA alters material properties of alpha-synuclein dense phase

While the irregular shape of tangles suggests a solid-like nature, neither confocal imaging nor svFCS can offer insights into the dense phase of aS when RNA is present. Specifically, FCS measurements show no correlation in any RNA condition. We speculate that the absence of correlation among fluorescence fluctuations stems from reduced liquid-like properties inside the tangles. The highly constrained environment of aS molecules within solid-like condensates hints toward the presence of a substantial immobile fraction, which makes FCS an unsuitable method to study dynamics. The presence of a significant immobile fraction and the irregular shape of the dense phase further support the hypothesis that RNA actively contributes to the transition towards solid-like aS condensates. For this reason, alternative approaches beyond FCS and related methods should be considered.

As a sensor of molecular crowding, fluorescence lifetime is a promising alternative for probing the different material properties of bio-molecular complexes (48). Notably, the SPAD array detector combined with the time-tagged detection allows the analysis of the photon-resolved dataset at both the micro-second scale requested by FCS and the sub-nanosecond scale requested by fluorescence lifetime. This allows us to correlate diffusion and fluorescence lifetime analysis on the same measurement (36, 38).

We measured the fluorescence lifetime of aS both in dilute and dense phases, with and without RNA. Generally, higher fluorescence lifetime values correspond to less packed, crowded nano-environments. In the dilute phase, we expect no variations in the fluorescence lifetime as svFCS measurements showed that it is primarily composed of monomeric protein. Indeed, except for the 5 ng µL RNA condition, the average fluorescence lifetime remains unchanged in the dilute phase (figure 4A). In the 5 ng µL RNA condition, we showed that the percentage of fast-diffusing monomers over the slow-diffusing nanoclusters is higher than that observed in the protein-only condition (figure 2E). This observation correlates with the increase in fluorescence lifetime (figure 3A).

**Fig 4.**
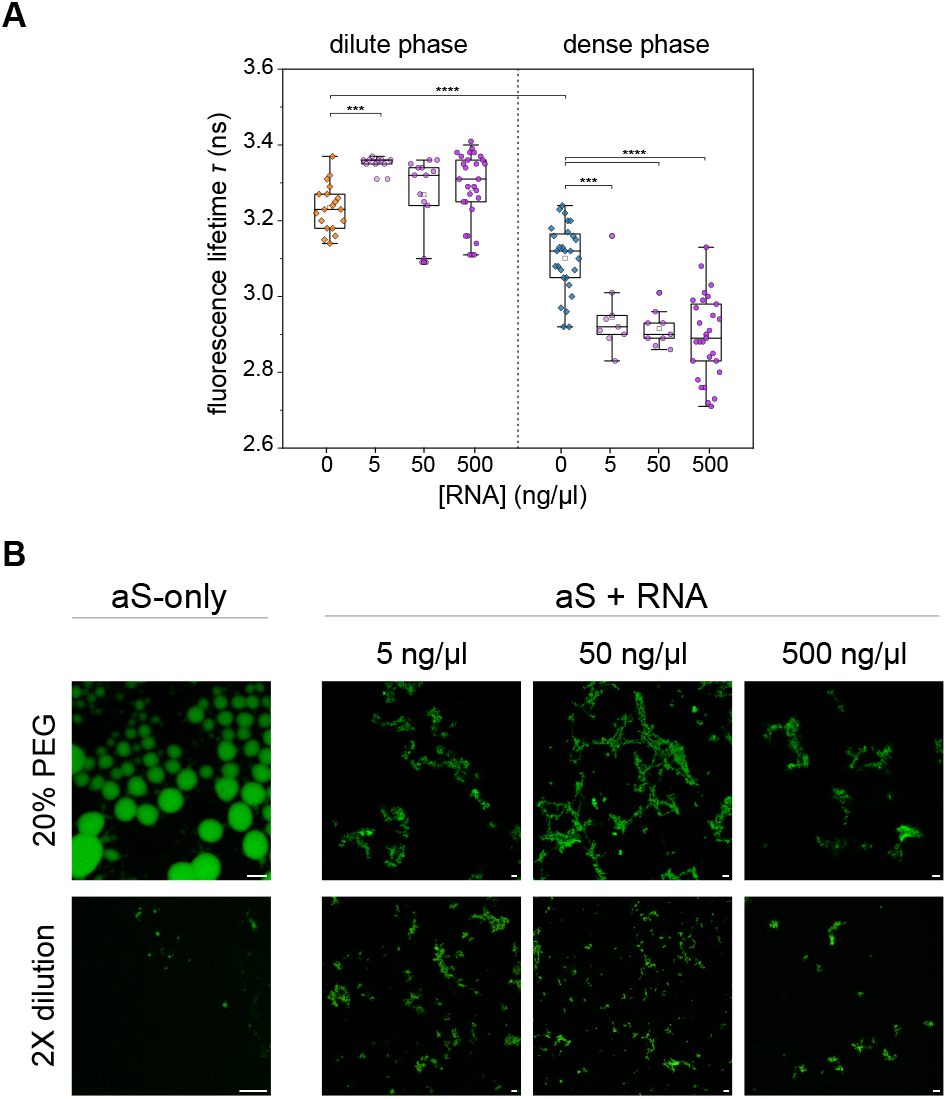
Fluorescence lifetime as a sensor for the condensation degree of aS in a phase-separated system. (A) Box plots of the fluorescence lifetime of aS-atto488 in the dilute (left) and dense (right) phase at different RNA concentrations. From left to right, *n* = 19, 10, 14, 29, 28, 9, 9, 29. One-way ANOVA and Kuskal-Wallis test followed by a Dunn’s test for the dilute phase (***: p-value *<* 0.0005, ****: p-value *<* 0.00005). (B) Confocal images of aS in LLPS buffer at different concentrations of RNA. Top is in LLPS buffer, bottom is after 2 times dilution with 20 µM PB.

Due to increased viscosity and molecular crowding, an overall reduction in the fluorescence lifetime values is expected in the dense phase compared to the dilute one. Indeed, lower average lifetime values were measured within condensates for all conditions. Surprisingly, a significative reduction in the fluorescence lifetime values is also observed between protein-only condensates and aS+RNA tangles. The reduction in the fluorescence lifetime is independent of the RNA concentration (figure 4A), suggesting that RNA increases the overall condensation/aggregation propensity of the protein irrespective of its concentration. This observation agrees well with the scenario in which RNA actively contributes to the liquid-to-solid transition of aS.

As further proof of our hypothesis, we tested the liquid-like properties and, in particular, the reversibility of the aS condensates in all four conditions. We diluted the 20% PEG buffer twice in PB. After 20 minutes upon dilution, we observed the almost complete dissolution of the aS-only condensates, while the aS+RNA tangles reduced their size but did not fully disappear (figure 4B). As the dilution reduces the crowding effect of the buffer (by effectively reducing the PEG concentration), if the condensates are purely liquid-like, they will dissolve; if they are partially solid-like, they will not.

## Discussion

Our study demonstrates the significant impact of RNA on the phase separation and aggregation of alpha-synuclein (aS). Using photon-resolved microscopy, we elucidate how the protein phase separates and how RNA modulates these processes at the nanoscale.

We initially studied aS phase separation in the absence of RNA, and we showed that nanoclusters form immediately and spontaneously in the dilute phase, even below the saturation concentration needed for macroscopic phase separation. This finding aligns with recent studies suggesting that phase separation can occur at much lower concentrations within cells than previously thought (18). The characterization of these clusters showed a higher degree of confinement compared to the compartmentalized component of the dense phase, likely due to the initial interaction dynamics during their formation. Once the nanoclusters grow into larger condensates, the intermolecular forces become more dynamic, resulting in a lower general degree of confinement.

We successively performed the same analysis in the presence of RNA, and we demonstrated that the nucleic acid promotes the liquid-to-solid transition of aS. This is evidenced by the reduction in the circularity of the dense phase, indicating the formation of irregular, more solid-like tangles. Fluorescence correlation spectroscopy measurements show no intensity fluctuations of fluorescent aS inside the dense phase with RNA, hinting at immobile molecules and confirming the transition to a solid-like state. Additionally, a significant decrease in fluorescence lifetime within aS+RNA tangles compared to aS-only condensates suggests higher molecular compaction and a conformational change of the protein in the presence of RNA.

The intrinsic flexibility of aS typically prevents stable interactions with RNA in its monomeric form. Computational predictions indicate a potential for RNA binding (49), which becomes biologically relevant during aggregation (23). As in aggregates, aS folds into structures that expose RNA-binding regions, facilitating interactions. This shift from non-binding to RNA-binding states underscores the dynamic nature of aS and its functional versatility in different aggregation states. Such interactions can modulate aS aggregation and phase separation processes, influencing the formation of pathological inclusions in neurodegenerative diseases.

In the dilute phase, aS exhibits two diffusing components: freely diffusing monomers and domain-confined nanoclusters. The presence of RNA does not alter the overall dynamics significantly but affects the monomer/nanocluster ratio and the confinement strength of aS within nanoclusters. At lower RNA concentrations, there is a reduction in nanocluster representation, indicating a potential negative impact of RNA on nanocluster formation. This highlights the nuanced role of RNA in modulating aS dynamics and suggests that the phase transition induced by RNA probably depends more on the type and sequence of the nucleic acid rather than its concentration.

In a more general context, the RNA structure can significantly impact the aggregation of proteins. As indicated by previous literature (50–52), certain RNA structures may promote or inhibit protein aggregation, depending on their interaction with the protein. We have observed that some RNA species are relevant for liquid-liquid phase separation, but it is also plausible that other species are crucial for the liquid-to-solid phase transition (53, 54). This distinction highlights the complexity of the role of RNA in protein aggregation and underscores the need for further research to delineate these effects.

Given the multiple spatial-temporal scales at which liquid-liquid phase separation (LLPS) processes occur, only a comprehensive approach integrating multiple quantitative methods can effectively investigate the underlying mechanisms. Our photon-resolved platform will be crucial for studying various LLPS processes and unveiling the mechanisms behind liquid-to-solid transitions (38). Furthermore, as this framework is compatible with living cells, it is also suitable for studying aS phase transitions in more complex environments where it is physiologically coupled with RNA. This understanding could ultimately facilitate the development of therapeutic strategies targeting the early stages of protein aggregation in neurodegenerative diseases.

## Materials and methods

### Alpha-synuclein purification and labeling

Wild type alphasynuclein (*α*-syn) and *α*-syn with the addition C141 (*α*-syn^C141^) were respectively purified as described in (23). Briefly, both proteins were recombinantly produced in *E. coli*. The pT7-7 aSyn C141 plasmid was a gift from Gabriele Kaminski Schierle (Addgene plasmid #108866; RRID: Addgene_108866). After lysis with boiling, ammonium sulfate precipitation, and dialysis, the proteins were subsequently purified via anion-exchange and size-exclusion chromatography with HiTrap™Q and HiLoad™16/600 Superdex™75 columns (Cytiva, MA, USA). The cysteine in position 141 was labeled *α*-syn^C141^ with atto-488 via a maleimide reaction according to the manufacturer’s instructions. Briefly, the dye is dissolved into water-free DMSO (Thermo Fisher, MA, USA) to a concentration of 10 mM. *α*-syn^C141^ dissolved in 20 mM phosphate buffer was incubated with 1.3 molar excess of atto-488-maleimide (ATTO-TEC GmbH, Germany) for 120 minutes at room temperature. After the incubation, the labeled protein is separated from the free atto-488 molecules via PD-10 desalting columns (Cytiva). The tagged protein was immediately checked for concentration and labeling ratio by UV-vis absorption spectroscopy (Nanodrop ND-1000, Thermo Scientific Technologies), then aliquoted, frozen, and kept at -80°C.

### Glass surface passivation

*µ*-Slide 8-well or 18-well (Ibidi GmHb, Germany) was passivated with methyl-poly-ethylene glycol silane 5000 kDa (mPEG-silane 5K, Sigma-Aldrich, JKA3037), as shown in (55). Briefly, multi-wells were cleaned by incubating them in 100% ethanol (Sigma-Aldrich) for 15 minutes, then sonicated for 10 minutes. After rinsing with ultra-pure water, multi-wells were incubated in 2% (v/v) Alconox detergent (Sigma-Aldrich) in ultra-pure water for 2 hours. Finally, *µ*-Slide 8-well were incubated in 1 g/l mPEG-silane 5K in 96 % ethanol and 0.02% (v/v) hydrochloric acid (Sigma-Aldrich) overnight, in agitation. After rinsing sequentially in ethanol and ultra-pure water, multiwells were incubated in 1% (v/v) Tween-20 (Sigma-Aldrich) in T50 buffer (10 mM Tris-HCl, 20 mM NaCl, pH 8) for 10 minutes, then rinsed in T50 buffer and ultra-pure water. Multi-wells were dried using nitrogen gas flow and stored at +4°C for a maximum of 1 week.

### In vitro liquid-liquid phase separation assay

The liquid-liquid phase separation assay was performed by diluting 50 µM full length aS and 2.5 µM (labeling ratio: 1:20) aS C141 atto-488-maleimide into the LLPS buffer (20% (wt/vol) polyethylene glycol 8000 kDa (PEG 8K, Sigma-Aldrich, Germany), 20 mM phosphate buffer (PB), 100 mM KCl and 5 mM MgCl_2_ (Sigma-Aldrich)). Only for *in vitro* phase separation assay of RNA or aS+RNA, dry powder of total yeast RNA (yRNA, Roche) was dissolved in 20 mM PB. After determining the concentration of the total yRNA mixture by UV-vis absorption spectroscopy (NanoDrop Microvolume Spectrophotometer, ThermoFisher Scientific), it was added to the LLPS buffer supplemented or not with aS. For the phase separation assay of RNA-only or aS+RNA, total yeast RNA was labeled with 200 nM SYTO™13 or SYTO™85 nucleic acid stain (ThermoFisher Scientific). The sample was immediately poured onto a previously passivated *µ*-Slide 8-well or 18-well (Ibidi GmHb, Germany) and imaged. All measurements were performed at room temperature.

### Optical architecture

#### Confocal imaging

Fluorescence images were acquired with a Leica SP5 (Leica Microsystems, Mannheim, Germany) inverted laser scanning microscope. Atto488/SYTO™13 and SYTO™85 were excited at 488 nm and 570 nm, respectively. We performed the experiment on a Leica 100x APO objective, NA = 1.40, oil dipping. We detected the fluorescence through a Leica HyD detector in the spectral region 510-550 nm and 580-620 nm, respectively for atto488/SYTO™13 and SYTO™85. Atto488 and SYTO™85 signals were acquired sequentially to avoid any cross-talk artifact. Phase contrast images and fluorescence images in figure 4B were acquired with a Nikon TiE inverted confocal microscope equipped with three excitation lasers (405, 488, and 561 nm) and three alkaline PMTs. We performed the experiment on a Nikon 100x APO objective, NA = 1.25, oil dipping. Atto488 was excited at 488 nm. The 488 nm laser was a CW diode semiconductor laser (Coherent).

#### Image analysis

The shape and circularity (defined as 4*π*(area/perimeter^2^)) analysis was performed with Fiji software as described in (56).

#### Fluorescence fluctuation spectroscopy

All the fluorescence fluctuation-based experiments have been performed on a custom laser-scanning microscope designed for photon-resolved microscopy with a SPAD array detector described in detail in (35, 36, 38, 57). The microscope uses a 5 × 5 SPAD array detector fabricated with a 0.16 µm BCD technology (34) and cooled to -15^°^C, resulting in a reduced background (57). Briefly, the microscope excites the sample with a triggerable 485 nm pulsed laser diode (LDH-D-C-485, PicoQuant, Berlin, Germany). We coupled the laser beam into a polarization-maintaining fiber before sending it to the microscope. The laser is controlled by a laser driver, which can be synchronized via an oscillator module (PDL 828-L “SEPIA II”, Picoquant). The oscillator provides the synchronization signal to feed into the time-tagging module to measure the photon-arrival times. The excitation beam is deflected by a pair of Galvo mirrors and focused on the sample using a 100 × /1.4 numerical aperture objective lens (Leica Microsystems, Wetzlar, Germany). The fluorescence signal is collected by the same objective lens, de-scanned by the two Galvo mirrors, and focused on the SPAD array detector. The measurements of monomeric alpha-synuclein have been acquired with the same optical system on a single-element SPAD detector (PDM-050-CTC-FC, Micro Photon Devices, Bolzano, Italy).

### Control and data-acquisition system for photon-resolved microscopy

The microscope is controlled by the BrightEyes microscope control suite (BrightEyes-MCS), which operates through a data acquisition board based on an FPGA (NI FPGA USB-7856R, National Instruments). The BrightEyes-MCS control unit drives the galvo mirrors and the axial piezo stage. It also records the signals collected by the SPAD array detector, all synchronized with the scanning beam system. The software, built on LabVIEW (National Instruments), utilizes the Carma application as its backbone (58). It provides a graphical user interface for controlling microscope acquisition parameters, registering digital signals from the SPADs, and visualizing recorded signals in real-time. When fluorescence lifetime measurements are combined with imaging or spectroscopy, we integrate the BrightEyes-TTM module (36) between the SPAD array detector and the FPGA-based control unit, as described in (36). In this setup, the BrightEyes-MCS control unit supplies the synchronization signals for the scanning process (i.e., pixel clock, line clock, and frame clock) to the BrightEyes-TTM. The BrightEyes-TTM also receives synchronization signals from the pulsed laser to measure photon arrival times relative to the fluorophore excitation event.

#### Experimental protocol

The detection volumes of the SPAD array detector have been calibrated by circular scanning FCS (57). Specifically, we measure the diffusion times of a sample of FluoroSpheres (YG, REF F8787, 2% solids, 20 nm diameter, actual size 27 nm, exc./em. 505/515 nm, Invitrogen, ThermoFisher, Waltham, MA, USA) diluted in ultrapure water 1 to 5000 and drop-casted on top of a glass coverslip. The fluorescence intensity was acquired for about 120 seconds and analyzed offline. All measurements were performed at room temperature. The detection volumes *ω*_0_ are 350, 370, 404, 420 nm for the SPAD array detector (respectively, from the smallest to the biggest *ω*_0_, figure S1) and 338 nm for the APD.

#### Fluorescence correlation calculation and analysis

We calculated the correlations directly on the lists of absolute photon times (59). For spot-variation analysis, the photon lists of all relevant SPAD channels were merged, and the correlations were calculated for each observation volume. The temporal data was split into chunks of 10 or 5 s, and for each chunk, the correlations were calculated. The individual correlation curves were visually inspected, and all curves without artifacts were averaged. For both single-point and circular scanning (60) FCS, the correlation curves were fitted with a 1-component or 2-component model, depending on the sample, assuming a Gaussian detection volume, as described in (57). A detailed description of the fitting functions can be found in the supporting information file. In the case of calibration measurements with fluorescent beads, one diffusing component model is used; in the case of alpha-synuclein measurements, typically, a two diffusing components model was necessary (it is stated in the manuscript when another model is used). The goodness of the fit was always checked by evaluating the reduced chi-squared and the Bayesian information criterion. For all FCS measurements acquired with the SPAD array detector, the correlation curves shown in the manuscript are related to the sum of the 25 SPAD channels. For the circular FCS measurements (60), the periodicity and radius of the scan movement were kept fixed while the autocorrelation amplitude, diffusion time, and focal spot size were fitted. This procedure was used for the fluorescent beads and allowed the different focal spot sizes to be calibrated. For the conventional FCS measurements, the focal spot size was fixed at the values found with circular scanning FCS, and the amplitude and diffusion times were fitted. The diffusion coefficient *D* can be calculated from the diffusion time *τ*_*D*_ and the focal spot size *ω*_0_ via 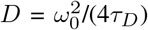. For the fluorescence lifetime measurements, we calibrated the microscope by measuring the instrument response function of the complete setup (microscope, detector, and BrightEyes-TTM) using a solution of atto-495 (fluorescence lifetime in H_2_O = 1 ns).

### Code availability

All the codes used for analyzing the photon time-tagging measurements (fluorescence lifetime fluctuation spectroscopy) have been deposited in the GitHub BrightEyes-TTM repository as a part of a larger openhardware/software project that aims at democratizing single-photon laser-scanning microscopy based on SPAD array detector (36), and it is available also on Zenodo (https://doi.org/10.5281/zenodo.7064910) (61).

### Statistical analysis

Statistical analysis was performed through OriginLab software, Version *2020* (OriginLab Corporation, Northampton, MA, USA). For each dataset, we performed a Shapiro-Wilk test to check the normality of the data distribution. When the hypothesis of normality was rejected, a non-parametric test (Mann-Whitney or Kruskal-Wallis tests) followed by a post hoc test (Dunn’s or Conover’s tests) was performed. When the hypothesis of normality was accepted, an ANOVA one-way parametric test followed by a Tukey test was performed. The exact p-values are reported in Table S1.

## AUTHOR CONTRIBUTIONS

G.V. and S.Z. conceived the idea. E.P., E.S., and G.V. developed the methodologies. E.Z., G.G.T., E.P. and G.V. supervised and coordinated the project. J.R. and S.Z. prepared the protein sample. S.Z. performed the experiments. E.P. and S.Z. analyzed the data. E.S. and G.V. designed and built, with the help of E.P., the custom laser-scanning microscope. E.S. designed and implemented the microscope control software. E.P. and E.S. designed the data analysis software. All authors discussed the results. S.Z., E.P., and J.R. wrote the manuscript with input from all authors.

## ACKNOWLEDGEMENTS

This project has received funding from: the Compagnia San Paolo, “Augmented fluorescence correlation spectroscopy with a novel SPAD array detector to observe complex biological processes in living cells”, Trapezio No. 71100 (E.P.); the European Research Council, “BrightEyes”, ERC-CoG No. 818699 (G.V.), “ASTRA”, ERC-SyG No. 855923 (G.G.T.); the European Innovation Council, “ivBM-4PAP”, Pathfinder No. 101098989 (G.G.T.); the NextGeneration EU PNRR MUR - M4C2 – Action 1.4 - Call “Potenziamento strutture di ricerca e creazione di “campioni nazionali di R&S” (CUP J33C22001130001), “National Center for Gene Therapy and Drugs based on RNA Technology”, No. CN00000041 (G.G.T. and G.V.). We thank all members of the Molecular Microscopy and Spectroscopy group and the RNA Systems Biology group at the Istituto Italiano di Tecnologia (IIT); Simone Civita and Matteo Mariangeli (Nanoscopy & NIC@IIT, IIT) for valuable discussions; Dr. Michele Oneto and Dr. Marco Scotto (Nikon Imaging Center, IIT) for support on the experiments; Prof. Alberto Tosi, Prof. Federica Villa, Dr. Mauro Buttafava (Politecnico di Milano), Dr. Marco Castello, and Dr. Simonluca Piazza (Genoa Instruments) for the realization of the single-photon-avalanche-diode detector array; all members of the RNA Initiative at the Istituto Italiano di Tecnologia, Prof. Stefano Gustincich (Non-coding RNAs and RNA-based therapeutics, IIT) and Dr. Francesco Nicassio (Genomics Science, IIT) for their contribution to the long-term vision of this project.

