## Supplementary material for "The role of RNA in the nanoscale organization of alpha-synuclein phase separation"

### Supplementary Note 1: Fluorescence correlation spectroscopy theory

Fluorescence Correlation Spectroscopy (FCS) is a microscopy method based on the single-molecule detection of fluorescence intensity fluctuations which can be used to describe the ensemble dynamics of a fluorescently tagged sample. FCS is part of a larger group of methods called fluorescence fluctuation spectroscopy. All these methods evaluate the fluorescence fluctuations to retrieve information about the dynamics and concentration of the sample. In FCS, the fluorescence intensity, which is acquired in a single point of the sample space over time, is autocorrelated in time at different time delays, the so-called lag times $\tau$, and the calculated autocorrelation function, $G(\tau)$, is fitted with an analytical model describing the type of dynamics of the sample inside a confined optical detection volume. For a detailed mathematical derivation of the analytical form of FCS, see (**?** ).

For the measurements on $\alpha$-synuclein, we fitted the autocorrelation curves with either a model for a single component freely diffusing in a 3D environment or a model for two components freely diffusing in 3D, with a confocal observation volume. For only one diffusing component in 3D:

$$G(\tau) = \frac{1}{\langle C \rangle V_{obs}} \frac{1}{1+\frac{\tau}{\tau_D}} \frac{1}{\sqrt{1+\frac{\tau}{\tau_D}(\omega_0/z_0)^2}}, \tag{S1}$$

where $\langle C \rangle$ is the average concentration of tagged molecules in the observation volume $V_{obs}$, $tau_D$ is the average diffusion time (defined as the time point at which the autocorrelation function reaches half of its maximum amplitude), and $\omega_0$ and $z_0$ are parameters describing the observation volume (the waist and the axial profile respectively).

In confocal microscopy, $V_{obs}$ is described by:

$$V_{obs} = (\pi/2)^{3/2}\omega_0^2 z_0, \tag{S2}$$

and needs to be calibrated before each measurement if the diffusion coefficient $D$ is the aim of the measurement. The diffusion coefficient can then be calculated from $\tau_D = \omega_0^2/4D$, and related to the size of the tagged molecule. If the tagged molecule is spherical, the radius $R$ can be estimated from the Stokes-Einstein equation:

$$D = \frac{K_B T}{6\pi\eta R}, \tag{S3}$$

with $K_B$ the Boltzmann constant, $T$ the temperature and $\eta$ the dynamic viscosity. The average number of molecules in the effective volume can be calculated from the average concentration as:

$$N = \langle C \rangle V_{eff}. \tag{S4}$$

In the case of multiple diffusing components, the autocorrelation function has multiple diffusing times. A model for two components diffusing in 3D in a confocal volume is described by:

$$G(\tau) = \frac{1}{N}(N_1 G_1(\tau) + N_2 G_2(\tau)), \tag{S5}$$

where $N$ is the average total number of tagged molecules in the observation volume, $N_1$ and $N_2$ are the percentage of number of molecules for respectively component 1 and component 2, and $G_1(\tau)$ and $G_2(\tau)$ represent the diffusion-related part of the autocorrelation function for each component. For example, for component one:

$$G_1(\tau) = \frac{1}{1+\frac{\tau}{\tau_{D1}}} \frac{1}{\sqrt{1+\frac{\tau}{\tau_{D1}}(\omega_0/z_0)^2}}, \tag{S6}$$

with $\tau_{D1}$ the diffusing time for component one.

For the calibration measurements circular scanning FCS has been performed. The fluorescence intensity is acquired over time by moving the laser beam in a circular orbit on the sample. The two parameters $\omega_0$ and $D$ can be decoupled by scanning the laser beam. In this case the autocorrelation function, $G(\rho,\tau)$ is a function of both spatial shifts $\rho$ and temporal lags $\tau$:

$$G(\rho,\tau) = \frac{1}{N\left(1+\frac{\tau}{\tau_D}\right)\sqrt{1+\frac{\tau}{\tau_D}}} \exp\left(-\frac{\rho^2}{\omega_0^2\left(1+\frac{\tau}{\tau_D}\right)}\right). \tag{S7}$$

In the case of a laser beam scanning in circles, $\rho(\tau)$ is written as:

$$\rho(\tau) = \sqrt{R^2+R^2-2R^2\cos(\gamma(\tau))} = \sqrt{2R^2\left(1-\cos\left(\frac{2\pi}{T}\tau\right)\right)}, \tag{S8}$$

DRAFT

where $R$ is here the radius of the scan pattern, $\gamma$ is the angle over the laser beam has traveled in a time interval $\tau$ and $T$ the time needed to scan a full circle. The parameter $\rho(\tau)$ corresponds to the distance between the two points under the angle $\gamma$. Combining Eq. S7 and S8 we obtain:

$$G(\rho,\tau) = \frac{1}{N\left(1+\frac{4D\tau}{\omega_0^2}\right)\sqrt{1+\frac{4D\tau}{z_0^2}}} \exp\left(-\frac{4R^2\sin^2\left(\frac{\pi}{T}\tau\right)}{\omega_0^2+4D\tau}\right). \tag{S9}$$

Since $D$ and $\omega_0$ now appear decoupled in the equation, they can be resolved together in a single experiment.

DRAFT

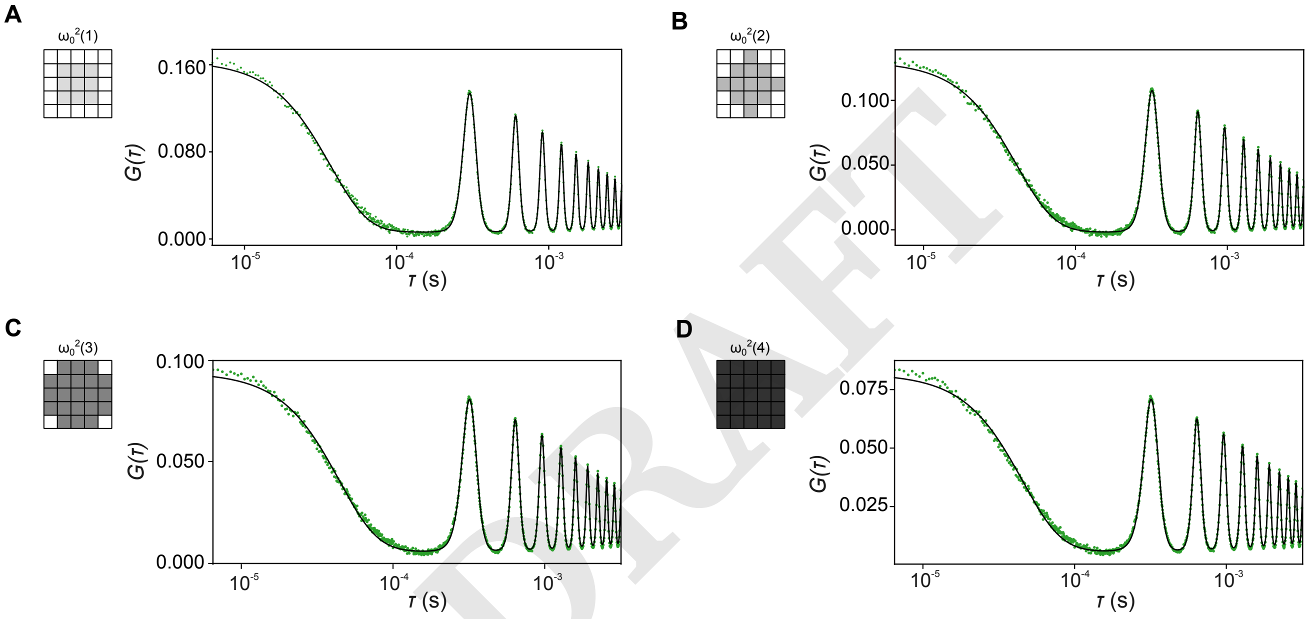


**Fig. S1. Determination of the lateral size of focal spots used for spot-variation FCS with circular scanning FCS on 20 nm fluorescent beads.** Correlation curves were fit with a freely diffusing model with $\omega_0$ and diffusion time $\tau_D$ as fit parameters. From the smallest to the biggest focal spot, the retrieved average $\omega_0$ values are 350, 370, 404, and 420 nm ($n$ = 10).

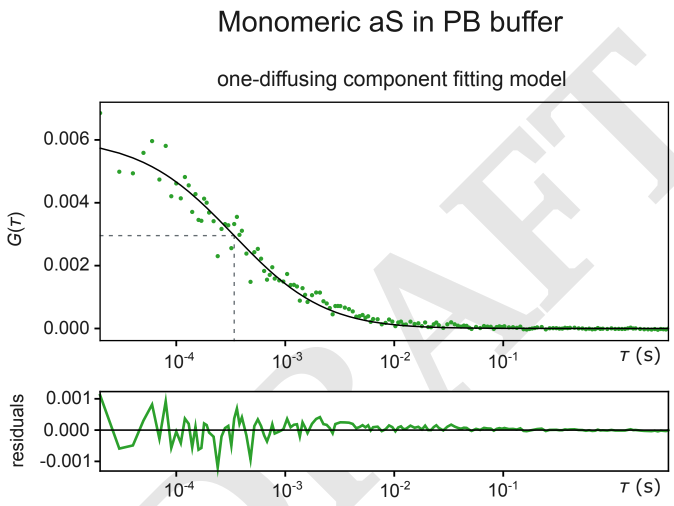


DRAFT

**Fig. S2. Representative autocorrelation curve *G(τ)* of 700 nM aS in 20 mM PB supplemented with 100 mM KCl and 5 mM $MgCl_2$.** The autocorrelation data was fitted with a one-diffusing component fitting model (solid line). Dashed lines indicate the corresponding diffusion time ($\tau_D$ = 0.37 ms).

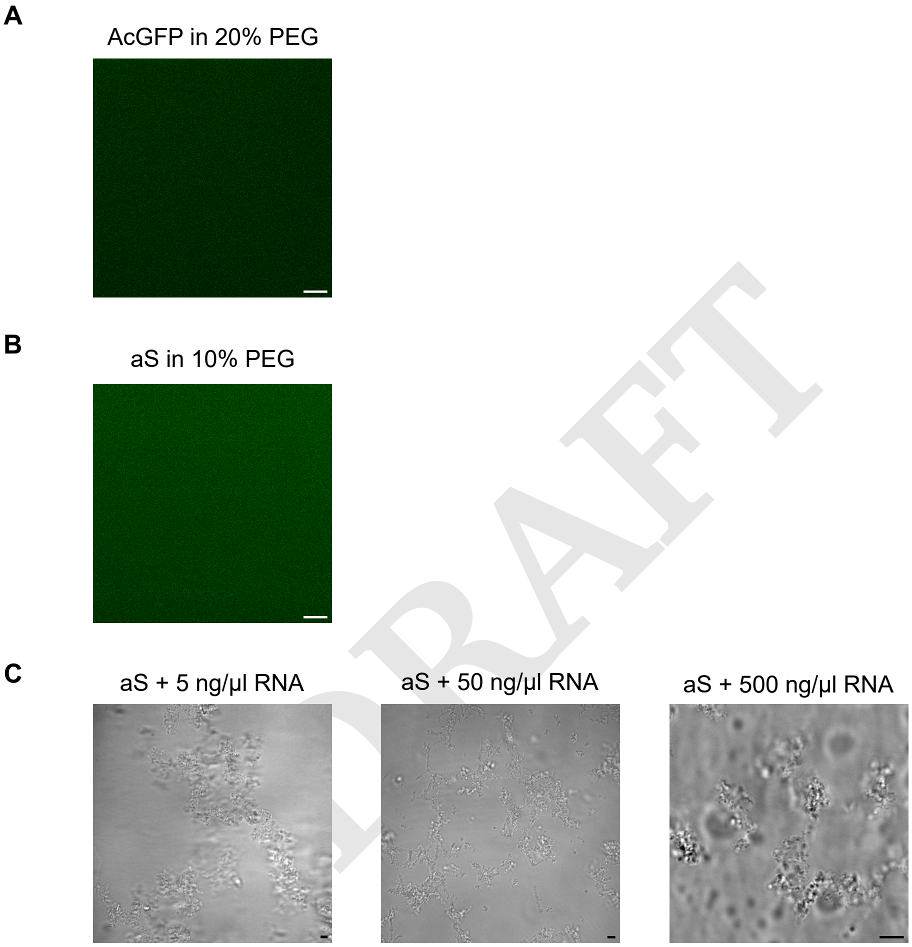


**Fig. S3. Confocal fluorescent and phase-contrast images.** (A, B) Confocal fluorescence images of (A) AcGFP in 20% PEG buffer and (B) labeled aS in 10% PEG buffer. In both cases, no micron-sized dense phase is appreciated. (C) Phase-contrast images of aS tangles upon addition of 5, 50 and 500 ng µL of RNA (from left to right). Scale bars: 5 µm.

DRAFT

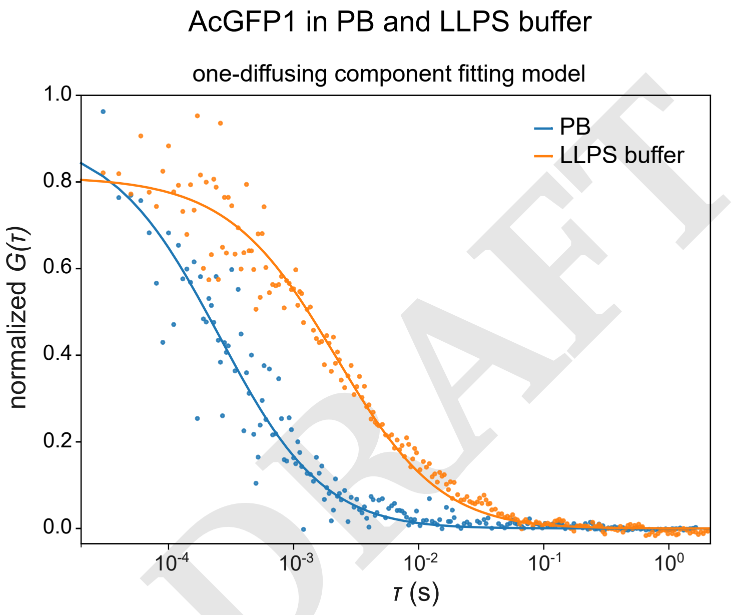


**Fig. S4. AcGFP1 as an estimator of the viscosity of LLPS buffer.** The figure shows a representative autocorrelation function $G(\tau)$ (scatter: data, lines: fitting model) of AcGFP1 (*Aequorea coerulescens* GFP) in 20 mM PB and LLPS buffer (20% PEG): $D_{app}$(PB) = 150 $\pm$ 13 µm$^2$ s, $D_{app}$(20% PEG) = 15 $\pm$ 1 µm$^2$ s. The comparison of both $D_{app}$ shows that the LLPS buffer is 10 times more viscous than PB.

DRAFT

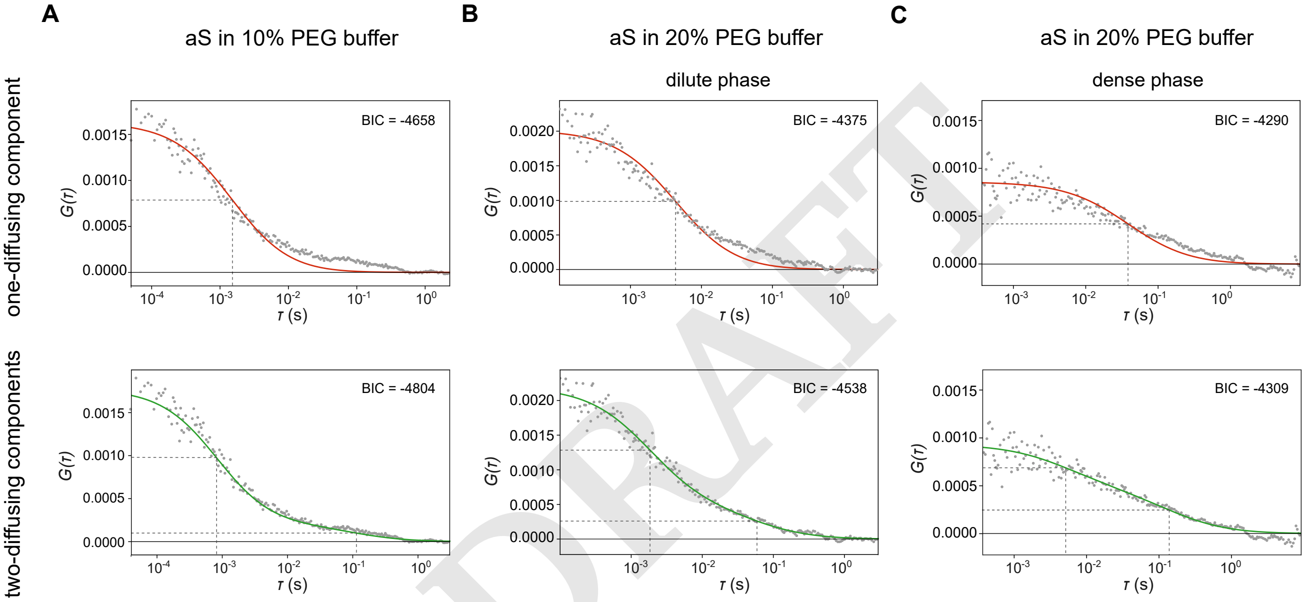


**Fig. S5. Representative autocorrelation functions $G(\tau)$ of aS in different LLPS buffers.** The autocorrelation functions (10% PEG for (A) and 20% for (B) and (C)) were fitted with one-diffusing component (red) or two-diffusing components (green) FCS model. The value of the Bayesian information criterion (BIC) is reported top-right for every $G(\tau)$ plot. The lowest BIC is generally correlated with a better-fitting model.

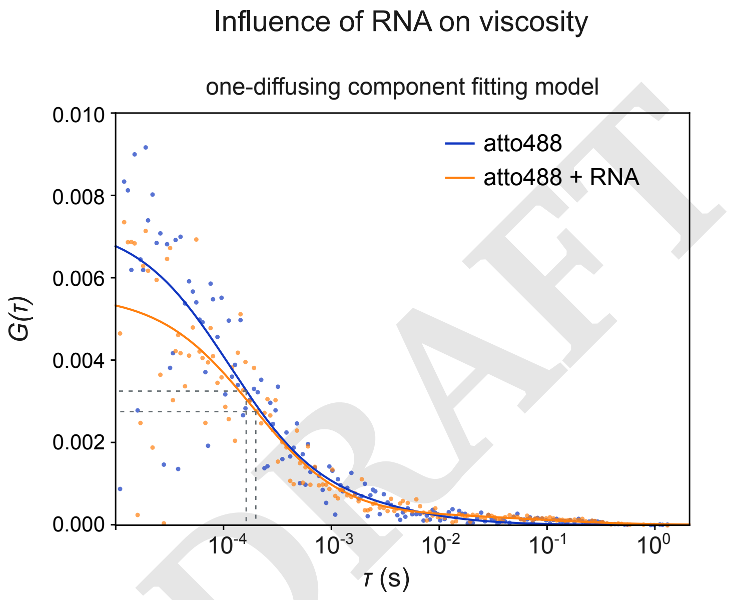


**Fig. S6. Evaluation of the influence of total yeast RNA on the viscosity of LLPS buffer.** The figure shows an example of the autocorrelation function of 100 nM atto488 in LLPS buffer (20% PEG) supplemented or not with 500 µM of RNA: $\tau_D$(atto488) = 0.15 $\pm$ 0.02 ms, $\tau_D$(atto488 + RNA) = 0.16 $\pm$ 0.02 ms. The diffusion time of atto488 does not change among the two conditions suggesting that RNA does not affect the viscosity of the LLPS buffer. FCS measurements were acquired with the APD.

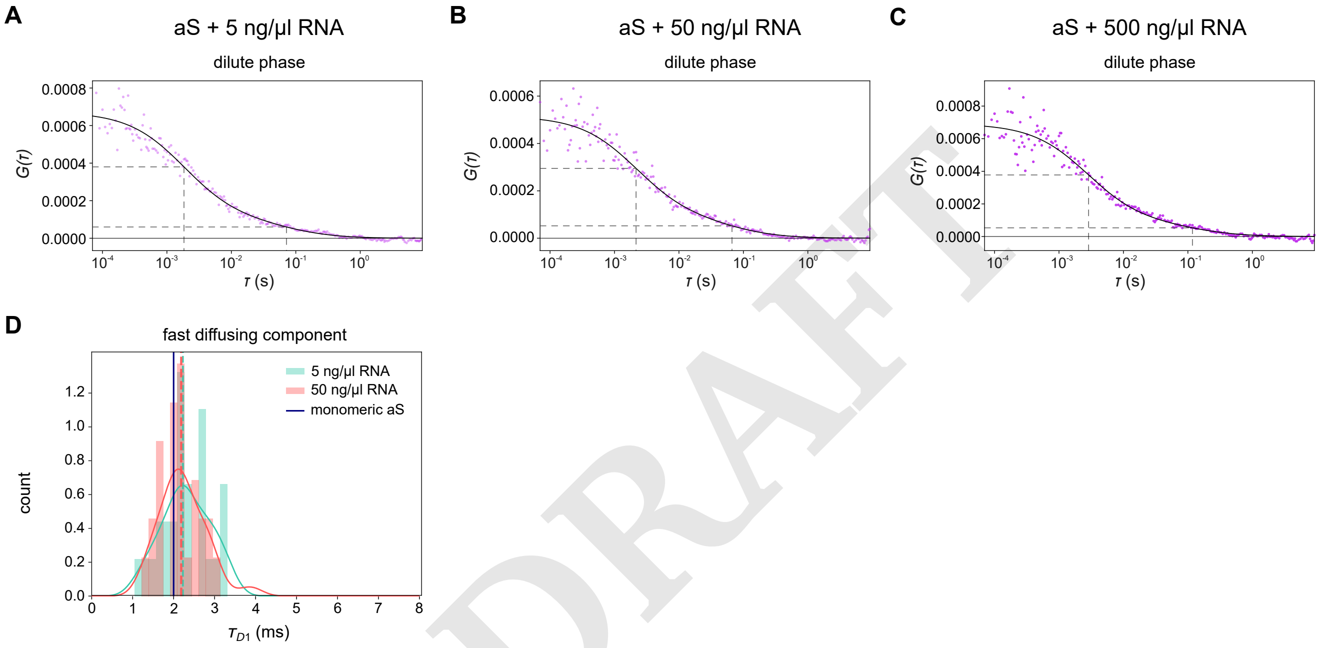


**Fig. S7. FCS measurements of aS dilute phase in presence of RNA.** (A, B, C) Representative autocorrelation function G($\tau$) of aS in the presence of 5 (A), 50 (B) and 500 (C) ng µL of total yeast RNA. Data have been fitted with a two-diffusing component model. Dashed lines indicate the diffusion time of both diffusing components; the faster component represents the freely diffusing monomeric aS, and the slower one represents the domain-confined nanoclusters. (D) Histogram of $\tau_{D1}$ of the fast component of aS in the presence of 5 and 50 ng µL RNA. Dashed lines represent the median (2.23 and 2.18 ms for respectively 5 and 50 ng µL RNA). The solid blue line represents the median of $\tau_{D1}$ associated with monomeric aS.

DRAFT

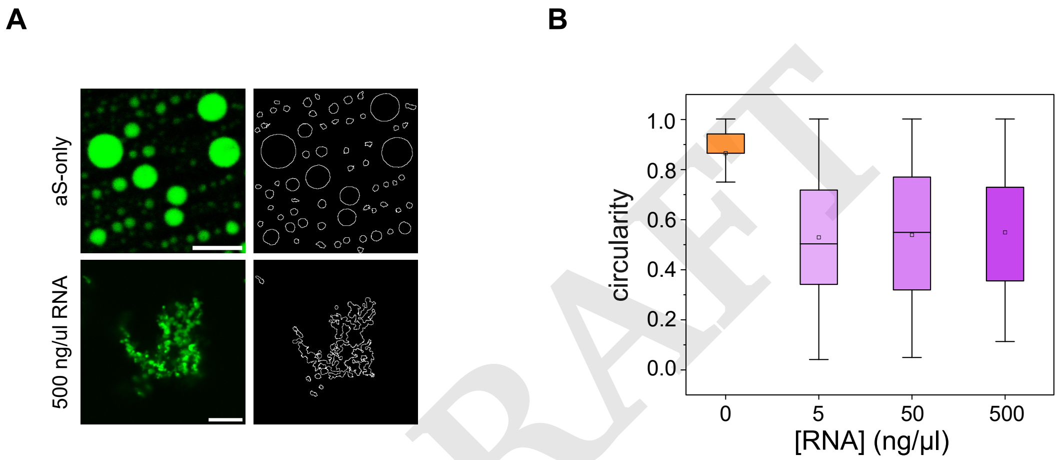


**Fig. S8. Analysis of the circularity of the dense phase of aS at different concentrations of total yeast RNA.** (A) Representative confocal images (left) and the outlines of the images segmented to measure the circularity of condensates (right). Scale bars: 5 µm. (B) Box plots of the circularity of the dense phase for aS alone (0) and in the presence of several RNA concentrations (5, 50, and 500 ng µL, from left to right). Circularity is defined as $\frac{4\pi * \text{area}}{\text{perimeter}^2}$. A value of 1 indicates a perfect circle; as the value deviates from 1, it indicates an elliptic shape. The outliers are not shown.

**Table S1.** Statistical analysis: P-values

| Sample | Test | P-value |
|---|---|---|
| %fast aS dilute - dense | Mann-Whitney | 2.5E-2 |
| $\tau_{D2}$ aS dilute - dense | Mann-Whitney | 4.7E-2 |
| $\tau_{D2}$ aS - 5 ng | Dunn's test | 2.8E-1 (n.s.) |
| $\tau_{D2}$ 5 ng - 50 ng | Dunn's test | 1 (n.s.) |
| $\tau_{D2}$ 50 ng - 500 ng | Dunn's test | 1.6E-1 (n.s.) |
| %(fast) aS - 5 ng | Dunn's test | 2.22E-4 |
| %(fast) aS - 50 ng | Dunn's test | 1.94E-5 |
| %(fast) aS - 500 ng | Dunn's test | 1.2E-1 (n.s.) |
| %(fast) 5 ng - 50 ng | Dunn's test | 1 (n.s.) |
| %(fast) 5 ng - 500 ng | Dunn's test | 6.6E-2 (n.s.) |
| %(fast) 50 ng - 500 ng | Dunn's test | 1.0E-2 |
| $\tau$ aS dilute - dense | Conover's test | 7.7E-6 |
| $\tau$ dilute aS - 5 ng | Conover's test | 4.7E-4 |
| $\tau$ dilute aS - 50 ng | Conover's test | 1 (n.s.) |
| $\tau$ dilute aS - 500 ng | Conover's test | 7.9E-2 (n.s) |
| $\tau$ dense aS - 5 ng | Conover's test | 1.5E-3 |
| $\tau$ dense aS - 50 ng | Conover's test | 5.7E-5 |
| $\tau$ dense aS - 500 ng | Conover's test | 7.3E-10 |

DRAFT
